# Reduced Capsaicin-Induced Mechanical Allodynia and Neuronal Responses in the DRG in the Presence of *Shp1* Overexpression

**DOI:** 10.1101/2024.01.23.576758

**Authors:** Robin Vroman, Shingo Ishihara, Spencer Fullam, Matthew J. Wood, Natalie S. Adamczyk, Nolan Lomeli, Fransiska Malfait, Anne-Marie Malfait, Rachel E. Miller, Adrienn Markovics

## Abstract

Transient Receptor Potential Vanilloid 1 (TRPV1) is a nonselective cation channel expressed by pain-sensing neurons and has been an attractive target for the development of drugs to treat pain. Recently, Src homology region 2 domain-containing phosphatase-1 (SHP-1) was shown to dephosphorylate TRPV1 in dorsal root ganglia (DRG) neurons, which was linked with alleviating different pain phenotypes. These previous studies were performed in male rodents only and did not directly investigate the role of SHP-1 in TRPV-1 mediated sensitization. Therefore, our goal was to determine the impact of *Shp1* overexpression on TRPV1-mediated neuronal responses and capsaicin-induced pain behavior in mice of both sexes. Twelve-week-old male and female mice overexpressing *Shp1* (Shp1-Tg) and their wild type (WT) littermates were used.

*Shp1* overexpression was confirmed in the DRG of Shp1-Tg mice by RNA *in situ* hybridization and RT-qPCR. *Trpv1* and *Shp1* were found to be co-expressed in DRG sensory neurons in both genotypes. Functionally, this overexpression resulted in lower magnitude intracellular calcium responses to 200 nM capsaicin stimulation in DRG cultures from Shp1-Tg mice compared to WTs. *In vivo*, we tested the effects of *Shp1* overexpression on capsaicin-induced pain through a model of capsaicin footpad injection. While capsaicin injection evoked nocifensive behavior (paw licking) and paw swelling in both genotypes and sexes, only WT mice developed mechanical allodynia after capsaicin injection. We observed similar level of TRPV1 protein expression in the DRG of both genotypes, however, a higher amount of tyrosine phosphorylated TRPV1 was detected in WT DRG. These experiments suggest that, while SHP-1 does not mediate the acute swelling and nocifensive behavior induced by capsaicin, it does mediate a protective effect against capsaicin-induced mechanical allodynia in both sexes. The protective effect of SHP-1 might be mediated by TRPV1 dephosphorylation in capsaicin-sensitive sensory neurons of the DRG.

## Introduction

Transient Receptor Potential Vanilloid 1 (TRPV1) is a nonselective cation channel expressed on a subset of unmyelinated C and medium diameter myelinated Aδ nociceptors [1]. TRPV1 has been an attractive drug target for the treatment of various types of pain, and both TRPV1 agonists and antagonists entered clinical trials. However, due to the physiological effects of TRPV1 on core body temperature and noxious heat sensation [2], the development of TRPV1 agonists and antagonists has been challenged by serious on-target adverse effects such as hyperthermia and impaired cutaneous noxious heat sensation [3, 4]. One of the major mechanisms of TRPV1 inhibition is dephosphorylation by protein phosphatases [5]. Interestingly, eliminating a serine phosphorylation site in TRPV1 by CRISPR/Cas9 editing reduced pain without affecting physiological TRPV1 functions in an inflammatory pain model [6], raising the possibility that modulating TRPV1 phosphorylation may offer an attractive strategy for targeting TRPV1 for analgesia.

Src homology region 2 domain-containing phosphatase 1 (SHP-1) is an intracellular protein tyrosine phosphatase expressed mainly in hematopoietic cells [7, 8]. It plays a negative regulatory role in immune cell signaling [9] and recently, its potential modulatory role in autoimmunity gained interest [10, 11]. A protective effect of SHP-1 has been proposed by others in different rodent models of pain. For example, SHP-1 has been shown to alleviate Complete Freund adjuvant (CFA)-induced thermal hyperalgesia [12]. It was also reported that SHP-1 dephosphorylates (i.e., deactivates) TRPV1 in dorsal root ganglia (DRG) and delays the development of cancer-induced bone pain in mice [13]. However, these previous studies were performed only in male rodents and did not directly investigate the role of SHP-1 in TRPV-1 mediated sensitization. Therefore, the objective of this study was to determine the impact of genetically enhanced SHP-1 expression on TRPV1-mediated neuronal responses and capsaicin-induced swelling and pain behavior in mice of both sexes. While selective SHP-1 inhibitors are commercially available, selective SHP-1 activators are lacking. Hence, SHP-1 transgenic mice (Shp1-Tg) with genetically enhanced SHP-1 expression were used to uncover the effect of increased SHP-1 activity.

## Materials and methods

### Animals

A total of 83 WT and 83 Shp1-Tg mice were used for the study. Animals were housed with food and water *ad libitum* and kept at a 12-hour light cycle. Twelve-week-old mice overexpressing *Shp1* (Shp1-Tg) on a BALB/c genetic background and their wild type (WT) littermates were used for the experiments. The generation of Shp1-Tg mice has been described elsewhere [10]. Mice were acclimatized to handling and to behavioral equipment for three days prior to the behavior studies. All animal experiments were approved by the Institutional Animal Care and Use Committee at Rush University Medical Center (IACUC #23-032).

### Drugs and chemicals

E-Capsaicin (04-621-00), Hank’s Balanced Salt Solution (HBSS, 14025-092), DMEM/F12 nutrient mixture (10-090-CV), and papain (NC9212788) were purchased from Fisher Scientific. Calbryte 520 AM (20650) was purchased from AAT Bioquest. Collagenase (C5138), DNase (10104159001), poly-L-lysine (P1274), laminin (L2020), Tween 80 (P4780), fetal bovine serum (FBS, SH30071.03, HyClone), N2 (17502-048), DMSO (D4540), penicillin-streptomycin (30-001-CI, MediaTech), ethanol and formaldehyde were purchased from Millipore-Sigma. SDS-polyacrylamide gels (10%, 4561034), PVDF membrane (1620177), blotting grade blocker (170-6404) and ECL Clarity Western blot substrate (170-5060) were purchased from Bio-Rad. BCA protein assay kit (23227) was purchased from ThermoFisher Scientific. Protein A/G Plus agarose beads (sc-2003) were purchased from Santa Cruz Biotechnology.

### Isolation of dorsal root ganglia

Briefly, mice were sacrificed, and the lumbar region of the spinal column was carefully dissected out as previously described [14]. The spine was cut in half in the sagittal plane one vertebra at a time and the spinal cord was removed. L3-5 DRGs were identified using a dissection microscope and excised with a micro dissection tool. These isolated DRGs were used for the following applications: 1) For RT-qPCR, bilateral L3-L5 DRGs of 6 naïve female WT and 5 naïve female Shp1-Tg mice were used. 2) For RNA *in situ* hybridization, bilateral L3-L5 DRGs of 3 naïve female WT and 3 naïve female Shp1-Tg mice were used. 3) For DRG culture and *in vitro* calcium imaging, bilateral L3-L5 DRGs of 3 female WT, 3 male WT, 3 female Shp1-Tg and 3 male Shp1-Tg mice were used.

### RT-qPCR

*Ptpn6* (the gene encoding SHP-1) gene expression was determined in the DRG of naive female mice (12 weeks old) by RT-qPCR. Briefly, total RNA was extracted with Direct-Zol RNA Mini-Prep Plus (Zymo Research, Irvine, CA) and reverse transcribed using the iScript Reverse Transcription Supermix for RT-qPCR (Bio-Rad, Hercules, CA). cDNA was amplified using SsoAdvanced Universal SYBR Green Supermix (Bio-Rad) in a CFX Connect Real-Time PCR Detection System (Bio-Rad). Measured ΔCq values were normalized to the actin b (*Actb*) gene. Relative expression was calculated using CFX Manager Software (Bio-Rad) and illustrated as ΔΔCq values relative to zero. *Ptpn6* primer sequences were as follows:

Forward: 5′-TCT CAG TCA GGG TGG ATG AT-3′

Reverse: 5′-CCT GCT GCT GCG TGT AAT A-3′

### DRG RNA *in situ* hybridization

Bilateral L3-L5 DRGs were collected from naïve female WT and Shp1-Tg mice (12 weeks old), fixed in 4% PFA, transferred to 30% sucrose solution for cryoprotection, embedded in OCT and cryo-sectioned onto slides at 12 μm. During sectioning slides were kept within the cryostat at −20 °C before storage at −80 °C. RNA *in situ* hybridization was performed using ACD BioTechne RNAscope Multiplex Fluorescent v2 Assay, described in detail elsewhere [15]. *Scn10a* (426011-C2 (probe for the gene encoding voltage-gated sodium channel Na_V_1.8)), *Trpv1* (313331-C3), *Ptnp6* (450081-C1) probes and ACD BioTechne DAPI were used. ACD Bio-Techne positive and negative control probes were conducted prior to start of work. Negative controls were included on every slide. All imaging was performed using a Fluoview FV10i confocal microscope at ×10 and ×60 magnification (Olympus Fluoview FV10-ASW Ver.04.02). Multiple planes of focus were captured, and the optimally focused image was chosen for processing and analysis. Laser intensity was used at ≤9.9% throughout. Images were processed and quantified using Fiji software v2.9.0. For mouse DRGs, 3 sections per mouse were quantified and averaged. First, the total number of neuronal cells was identified using both the nuclei staining with DAPI and the phase contrast channel. Each cell was then assessed for the expression of each probe, and labeled as single, double, triple expression, or no expression. Positive signal was determined when 2 or more positive ‘dots’ per cell were found.

### DRG culture and calcium imaging

Bilateral L3-L5 DRGs were collected from 12-week-old naïve Shp1-Tg and WT mice and pooled for enzymatic digestion using collagenase type IV (2 mg/ml), papain (26 U/ml), and DNase (1 mg/ml), as described [16]. Dissociated DRG cells were plated on poly-L-lysine and laminin-coated glass coverslips in 6-well plates and cultured in F12 medium supplemented with 1 × N2, 0.5% FBS, penicillin and streptomycin (100 µg/ml and 100 U/ml), at 37 °C and 5% CO_2_. Cultured DRG neurons were used for *in vitro* calcium imaging in two independent experiments (n=2-3 coverslips per genotype/sex/experiment). Neuronal responses were evaluated on four-day old DRG cultures by *in vitro* calcium imaging using Calbryte 520 AM calcium indicator dye, using a standard protocol [17]. Following 20 minutes of Calbryte loading, cells were rinsed and left untouched for 30 minutes to recover in a balanced salt solution [NaCl (140 mM), HEPES (10 mM), CaCl_2_ (2 mM), MgCl_2_ (1 mM), Glucose (10 mM), KCl (5 mM)] before being placed in an imaging chamber with a continuous flow system. Baseline neuronal activity was recorded for 5 minutes, followed by the application of vehicle (0.4 % DMSO) or 200 nM capsaicin to the imaging chamber. Neuronal responses were visualized under an inverted microscope (Zeiss Axio Observer D1). DRG cultures from WT and Shp1-Tg mice were evaluated on the same day. Neuronal response was quantified as the change in fluorescent intensity at a given time point compared to baseline fluorescence (ΔF/F_0_) using Image J software. Positive neuronal response was defined as maximum ΔF/F_0_ above 0.75.

### Capsaicin-induced acute pain behavior

Twelve- to fourteen-week-old naïve WT and Shp1-Tg mice of both sexes (n=31 female WT, n=30 female Shp1-Tg, n=27 male WT, n=29 male Shp1-Tg) were habituated to a plexiglass chamber and to the von Frey apparatus for three consecutive days. Mice were injected with 12 µg capsaicin or vehicle (80% saline, 10% ethanol, 10% Tween80) in the left footpad (10-20 µL) using a Hamilton syringe (n=14-17/treatment group). Capsaicin-induced paw licking, as a measure of nocifensive behavior, was recorded for 5 minutes in a transparent plexiglass chamber immediately after footpad injection and quantified as the time spent with paw licking during the first 5 minutes after injection [18]. After that, mice were returned to their home cage and allowed to rest until 4 hours after footpad injection, when mechanical allodynia was determined using calibrated von Frey filaments and the up-down staircase method [16]. Briefly, mice were placed on a perforated metal grid with plexiglass cubicles. A set of six von Frey fibers (Stoelting Touch Test Sensory Evaluator Kit) were applied to the plantar surface of the hind paw until they bowed. The force that resulted 50% paw withdrawal threshold was calculated as previously described [19]. Five hours after footpad injection, the thickness of the injected paw was determined using a caliper. The investigator was not blinded to the *in vivo* experimental groups, due to the obvious swelling and redness of the paw following capsaicin footpad injection.

### Immunoprecipitation and Western blot

For immunoprecipitation, ipsilateral L3-L5 DRGs of 10 female WT and 10 female Shp1-Tg mice were used (12 weeks of age) in two independent experiments. WT and Shp1-Tg female mice were injected with 12 µg capsaicin in the left footpad (n=5/genotype/experiment). Ipsilateral L3-L5 DRGs were harvested four hours after footpad injection and a whole cell lysate was prepared using RIPA buffer containing protease and phosphatase inhibitors. The protein concentrations were determined using a BCA Protein Assay kit as described previously [10]. For immunoprecipitation, 300 µg cell lysates were incubated with anti-phospho-tyrosine antibody (#8954, Cell Signaling Technology) at 4 °C overnight. Protein A/G Plus agarose beads were added to the lysate-antibody mixture and incubated for 4 hours. Samples were washed and eluted in reducing sample buffer, followed by boiling for 5 minutes. As input, 10 µg whole cell lysates were used. Proteins were separated on 10% SDS-polyacrylamide gels, blotted onto PVDF membranes, then blocked with Tris-buffered saline + 0.05% Tween-20 containing 3% blotting-grade blocker. The membrane was probed with anti-TRPV1 antibody (NBP1-97417, Novus Biologicals) at 4 °C overnight, followed by goat anti-rabbit HRP-conjugated secondary antibody (31466, ThermoFisher Scientific). Clarity ECL Western Blotting Substrate was used to generate chemiluminescence signals, which were detected by X-ray films.

### Statistical analysis

Results are presented as the mean ± SEM. RT-qPCR data were analyzed with Mann Whitney test. RNA *in situ* hybridization data and area under the curve values from DRG neuronal responses were compared with unpaired Student’s t-test. Data obtained *in vivo* (time spent with paw licking, paw thickness and mechanical allodynia) were analyzed with Kruskal-Wallis test followed by Dunn’s multiple comparison. Paw withdrawal threshold data were logarithm transformed prior to statistical analysis. Analyses were carried out using GraphPad Prism v9.3.1 software. A p-value of less than 0.05 was considered statistically significant.

## Results

### *Ptpn6* is overexpressed in the DRG of Shp1-Tg mice and co-expressed with *Trpv1* in nociceptors

First, we aimed to confirm SHP-1 (encoded by the *Ptpn6* gene) overexpression in the DRG of Shp1-Tg mice by RT-qPCR. We found significantly higher *Ptpn6* expression in the DRG of Shp1-Tg mice compared to WT mice **(Figure 1A)**.

**Figure 1.**
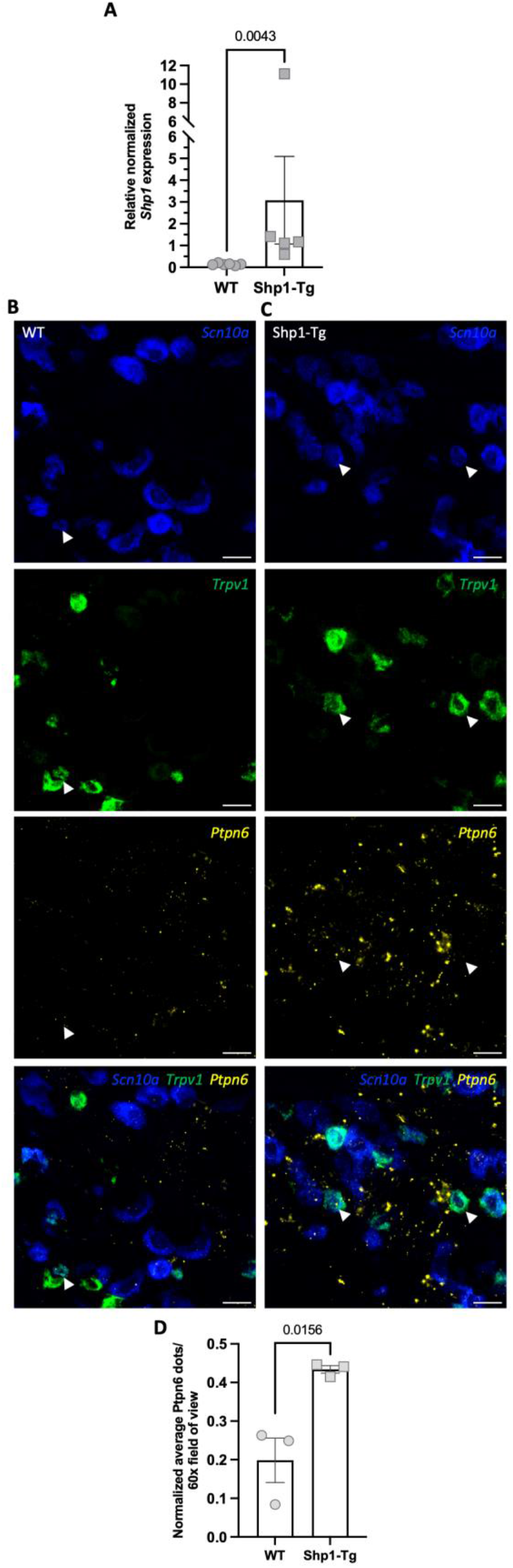
*Shp1* expression in the DRG. **A**: *Shp1 (Ptpn6)* gene expression in the DRG of WT and Shp1-Tg mice, quantified by RT-qPCR (naïve females, n=5 WT, n=6 Shp1-Tg, Mann Whitney test). **B-C**: Representative RNA *in situ* hybridization images of the DRG of n=3 WT (column B) and n=3 Shp1-Tg mice (column C). **Top to bottom**: blue-*Scn10a*, green-*Trpv1,* yellow-*Ptpn6*. White arrowheads are pointing to positive signal of the respective staining. Overlay: arrowhead is pointing to cells positive to *Ptpn6, Trpv1 and Scn10a*. Scale bar: 25 µm. **D**: *Ptpn6* expression quantified based on fluorescence by RNA *in situ* hybridization (naïve female, n=3/genotype, unpaired t-test). Data are expressed as mean ± SEM.

Next, we sought to determine if *Ptpn6* and *Trpv1* are both expressed in the nociceptors of the L3-L5 DRG, allowing for the potential modulation of TRPV1 function by SHP-1. The expression of *Trpv1* (*Trpv1* probe, green) and *Ptpn6* (*Ptpn6* probe, yellow) were visualized in nociceptive sensory neurons (*Scn10a* probe encoding Na_V_1.8, dark blue) of the DRG in both genotypes by RNA *in situ* hybridization. We found that some of the *Scn10a* positive sensory neurons were also positive for the *Ptpn6* and *Trpv1* probes, indicating co-expression of *Ptpn6* and *Trpv1* in the DRG sensory neurons of both genotypes (representative images of DRGs from n=3 naïve WT mice and n=3 naïve Shp1-Tg mice are shown in **Figure 1B-C).** We observed pronounced *Ptpn6* expression in *Scn10a* negative cells as well **(Figure 1B-C, overlay)**, which might be due to abundant *Ptpn6* expression by hematopoietic cells [8] that can be present in the DRG. Finally, overall *Ptpn6* expression in the DRGs was quantified by normalizing the average number of *Ptpn6* dots per 60X field of view. *Ptpn6* was found to be elevated in the DRG of Shp1-Tg mice compared to WT mice **(Figure 1D)**, confirming the results observed by RT-qPCR.

### Reduced capsaicin-induced neuronal responses in the DRG of Shp1-Tg mice

To evaluate the capsaicin-induced neuronal activation in the L3-L5 DRGs of WT and Shp1-Tg mice, neuronal responses were recorded *in vitro* in cultured DRG neurons from both genotypes and sexes, using calcium imaging (**Figure 2**). Representative neuronal responses are illustrated in **Figure 2A-D** (the number of responding neurons to 200 nM capsaicin with peak ΔF/F_0_ values above the cutoff value was 42 out of 108 for cultures of female WT, 20 out of 65 for female Shp1-Tg, 13 out of 60 for male WT and 3 out of 28 for male Shp1-Tg culture, respectively). Because we did not detect differences in responses between sexes, we combined male and female data and performed area under the curve analysis of all responding neurons. This analysis indicated significantly lower responses in the DRG cultures of Shp1-Tg mice compared to WTs (p=0.023) **(Figure 2E)**. Vehicle (DMSO) administration did not elicit neuronal responses (not shown).

**Figure 2.**
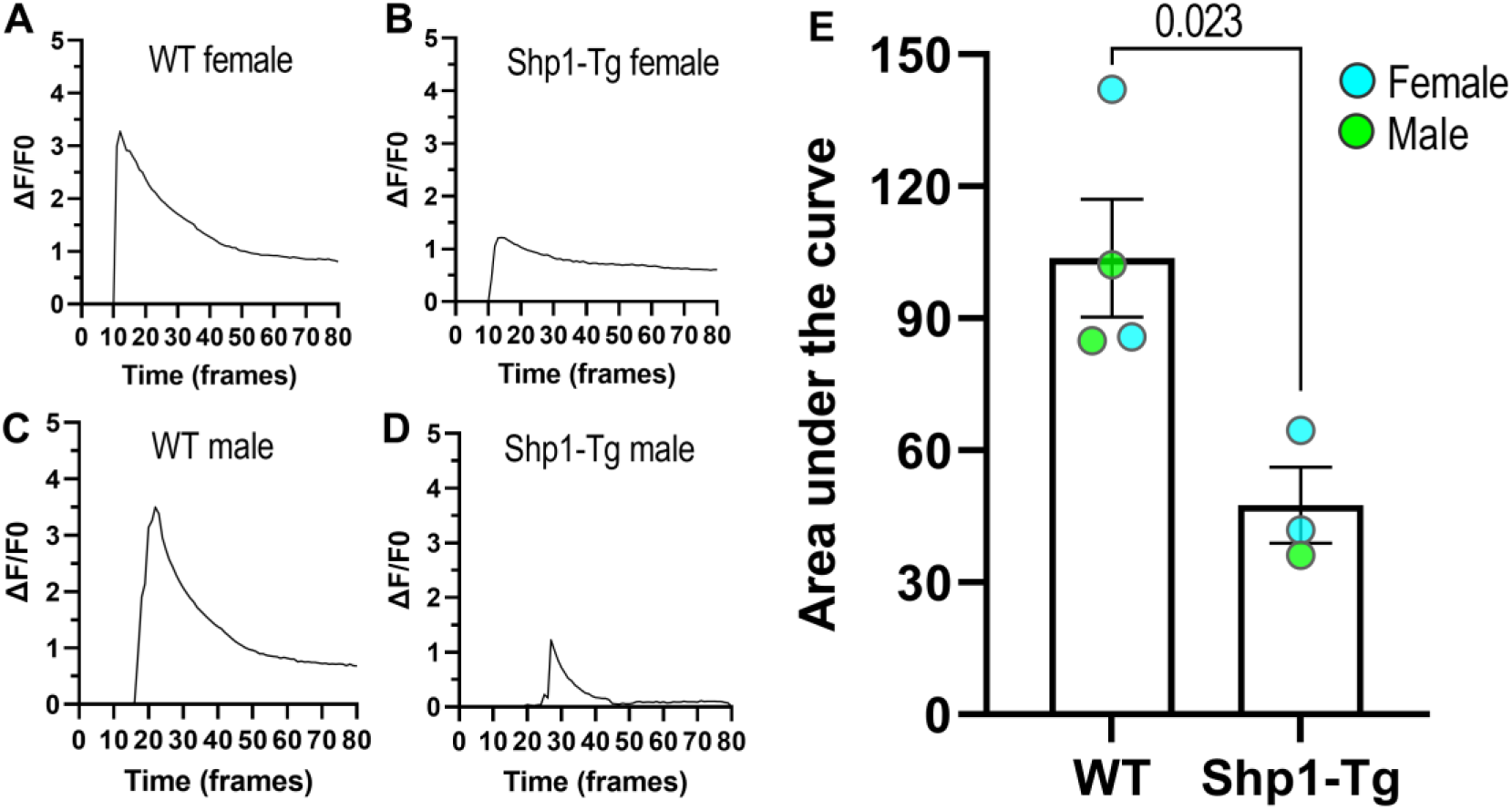
Capsaicin induced neuronal responses of the DRG *in vitro*. Capsaicin-evoked responses of cultured DRG neurons were analyzed for WT and Shp1-Tg cultures of both sexes. A, B, C, D: Representative individual neuronal responses to 200 nM capsaicin stimulation. E: Area under the curve of neuronal responses from WT and Shp1-Tg DRG cultures (mean ± SEM, n=3-4/group, Student’s t-test, two independent experiments/genotype).

### Genetically enhanced SHP-1 expression did not affect capsaicin-induced paw licking and paw swelling

Twelve- to fourteen-week-old female and male WT and Shp1-Tg mice were used for behavioral testing. As a measure of capsaicin-induced acute nocifensive behavior, we monitored the time spent with paw licking in the first five minutes after capsaicin or vehicle footpad injection. WT and Shp1-Tg mice of both sexes spent significantly more time with paw licking in response to capsaicin than to vehicle **(Figure 3A-B)**. There was no significant difference in capsaicin-evoked paw licking between WT and Shp1-Tg mice. Male mice of both genotypes appeared to spend less time with paw licking in response to capsaicin compared to females, however the difference was not statistically significant (WT capsaicin female vs WT capsaicin male: p=0.278; Shp1-Tg capsaicin female vs Shp1-Tg capsaicin male: p=0.067). Interestingly, while WT mice showed no paw licking after vehicle injection, some of the Shp1-Tg mice - especially females-displayed a short paw licking response after vehicle (**Figure 3A-B**), however this was not statistically significant between genotypes. The thickness of the injected paw was determined 5 hours after footpad injection. Capsaicin injection led to paw swelling indicated by increased paw thickness compared to vehicle in both genotypes **(Figure 3C-D)**. No significant difference was detected in the effect of capsaicin on paw thickness between the genotypes.

**Figure 3.**
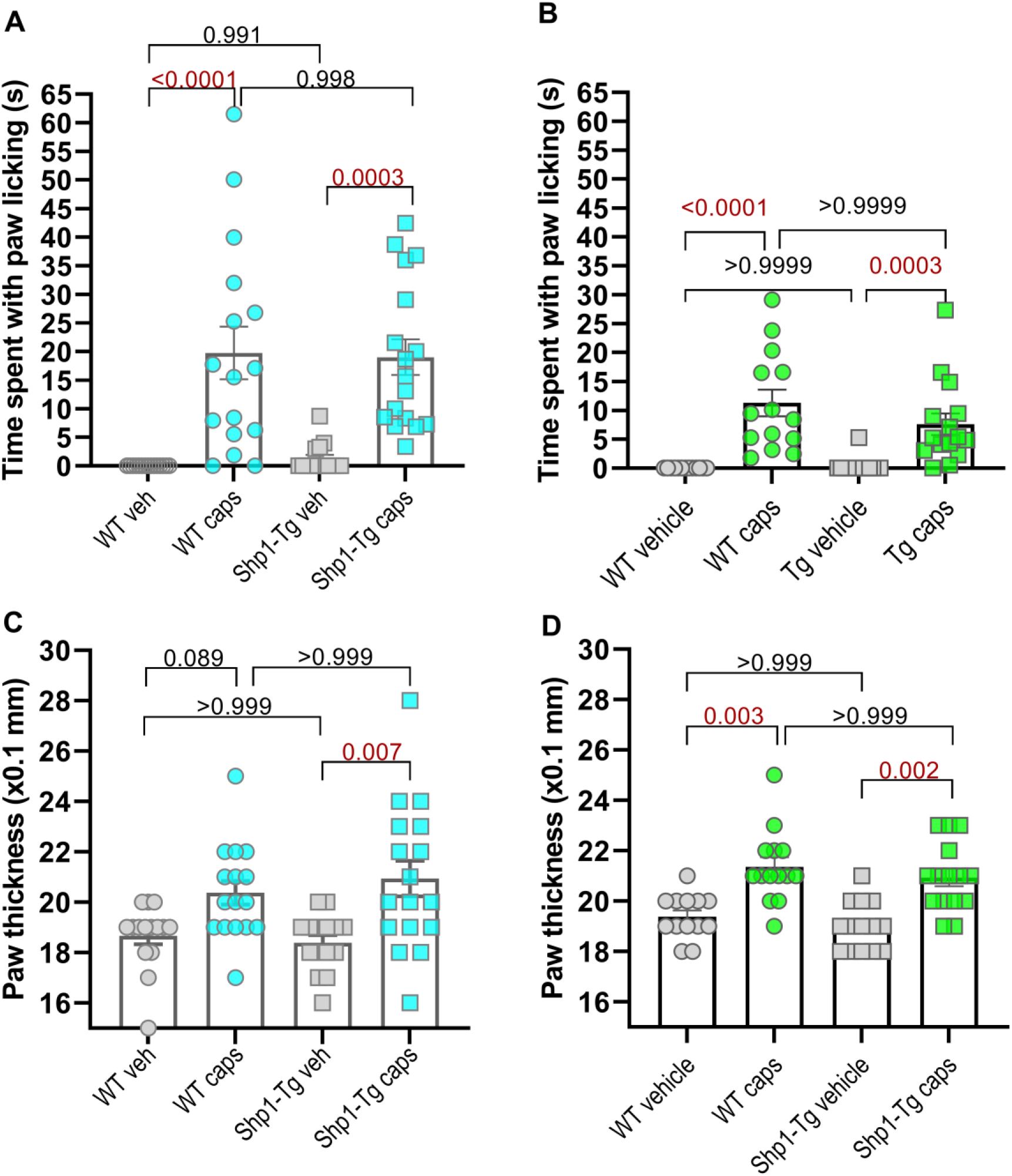
Capsaicin-induced acute nocifensive behavior and paw swelling in WT and Shp1-Tg mice. A-B: Time spent with paw licking in female (A) and male (B) mice (n=14-17/group, mean ± SEM, Kruskal-Wallis test and Dunn’s multiple comparison). C-D: Paw thickness in female (C) and male (D) mice after capsaicin or vehicle footpad injection (n=14-17/group, mean ± SEM, Kruskal-Wallis test and Dunn’s multiple comparison). p-values lower than 0.05 are indicated in red font.

### Shp1-Tg mice show reduced capsaicin-induced mechanical allodynia

As a measure of mechanical allodynia, we determined the paw withdrawal threshold with von Frey filaments 4 hours after capsaicin or vehicle footpad injections. Capsaicin induced a significant drop in the 50% withdrawal threshold in both female and male WT mice compared to vehicle (**Figure 4A-B**). Remarkably, neither female nor male Shp1-Tg mice exhibited significant mechanical allodynia induced by capsaicin compared to vehicle (**Figures 4A-B**).

**Figure 4.**
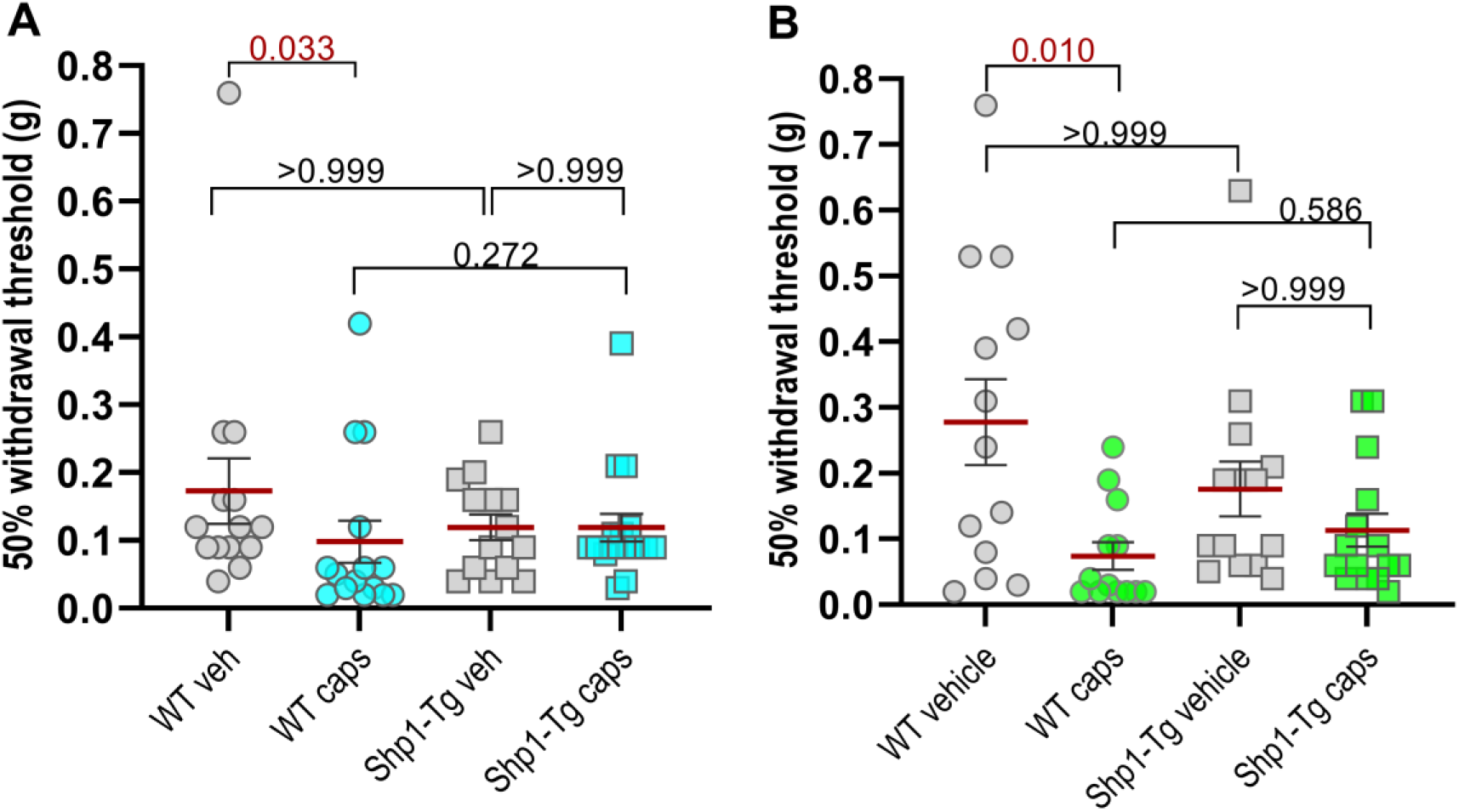
Capsaicin-induced mechanical allodynia in WT and Shp1-Tg mice. Paw withdrawal threshold after capsaicin or vehicle administration in female (A) and male (B) mice (n=14-17/group, mean ± SEM, Kruskal-Wallis test and Dunn’s multiple comparison). p values lower than 0.05 are indicated in red font.

### Reduced tyrosine phosphorylation of TRPV1 in the DRG of Shp1-Tg mice

To further investigate the observed protective effect on mechanical allodynia, we compared the tyrosine phosphorylation status of TRPV1 in the L3-L5 DRG of a separate cohort of WT and Shp1-Tg mice 4 hours after injection with capsaicin **(Figure 5)**. Total tyrosine phosphorylated proteins were immunoprecipitated from tissue lysates of both genotypes and immunoblotted with an anti-TRPV1 antibody. 10 µg tissue lysate without immunoprecipitation from both genotypes served as input for total TRPV1 immunoblotting. We detected a similar amount of total TRPV1 in WT and Shp1-Tg DRG lysates (lane 1 and 2). In contrast, while we observed a band according to tyrosine phosphorylated TRPV1 in the lysate of WT mice (lane 3), we did not detect tyrosine phosphorylated TRPV1 in the lysate of Shp1-Tg mice (lane 4).

**Figure 5.**
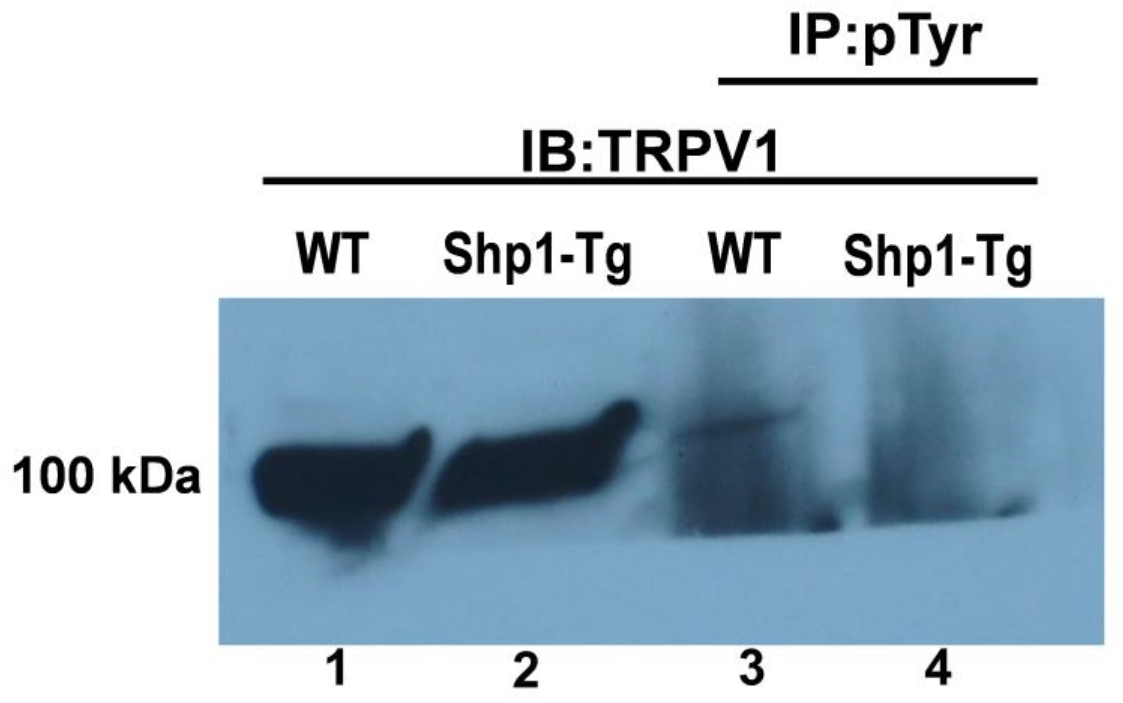
Tyrosine phosphorylation status of TRPV1 in the DRG. Lane 1-2: Immunoblot of DRG protein extracts from female WT and Shp1-Tg mice, probed with anti-TRPV1. Lane 3-4: immunoblot of immunoprecipitated tyrosine phosphorylated proteins from DRG protein extracts of WT and Shp1-Tg mice, probed with anti-TRPV1. Representative of n=2 experiments.

## Discussion

The aim of this study was to investigate the effect of genetically enhanced *Shp1* expression on TRPV1-mediated neuronal responses to capsaicin in the DRG as well as on TRPV1-mediated swelling and pain behavior using WT and Shp1-Tg mice. Here we confirm that *Shp1* and *Trpv1* are both expressed in the nociceptors of the L3-L5 DRG. In addition, we detected reduced neuronal responses to capsaicin in the DRGs of Shp1-Tg mice. Capsaicin footpad injection led to nocifensive behavior and swelling in both genotypes and sexes, while reduced capsaicin-evoked mechanical allodynia was observed in the presence of genetically enhanced *Shp1* expression. Furthermore, reduced tyrosine phosphorylated TRPV1 was detected in the DRG of Shp1-Tg mice.

Our results support previous findings that demonstrated a protective role of SHP-1 in other types of inflammatory pain models. For example, SHP-1 alleviated CFA-induced inflammatory pain in rats [12] by counteracting the tyrosine phosphorylation of TRPV1 by Src (a non-receptor protein tyrosine kinase) in the L4-L5 DRG of rats. SHP-1-TRPV1 interaction in DRG neurons has also been reported by Liu *et al*. [13] who found that Programmed Death Ligand-1 (PD-L1) diminished bone cancer-induced pain in mice through suppressing TRPV1 function, and this effect was mediated by SHP-1 in the DRG. However, when the role of SHP-1 in the central nervous system was examined, SHP-1 was found to mediate pain through effects independent of TRPV1 [20], and *Shp1* knockdown by siRNA in the spinal cord alleviated CFA-induced inflammatory pain [21]. These contradictory data on the role of SHP-1 in pain can be explained by TRPV1-independent mechanisms and different levels of the pain pathway affected. Our findings indicate reduced mechanical allodynia after capsaicin injection in Shp1-Tg mice, outlining a protective role of SHP-1 in TRPV1-mediated sensitization.

Phosphorylation plays an important role in the modulation of TRPV1 activity. Src can phosphorylate TRPV1 at Tyr200, leading to TRPV1 trafficking to the plasma membrane and an increase of TRPV1 responses in the DRG [22, 23]. Interestingly, many of the substrates of Src can be subject to dephosphorylation by SHP-1 [24]. SHP-1 has been shown to abolish increased TRPV1 tyrosine phosphorylation by Src in cultured HEK-293 cells [23]. SHP-1 might also counteract NGF-induced TRPV1 sensitization, as it was reported that NGF binding to its receptor, TrkA, initiates a signaling cascade in which Src kinase phosphorylates TRPV1 in a single tyrosine residue, promoting its traffic to the plasma membrane and contributing to NGF-induced sensitization [23].

We found a similar total TRPV1 protein expression in the L3-L5 DRG of both WT and Shp1-Tg mice, while there was much less tyrosine phosphorylated TRPV1 in the DRG of capsaicin-injected Shp1-Tg mice. Therefore, we hypothesize that the reduced mechanical allodynia is conveyed by the enhanced dephosphorylation of TRPV1 by SHP-1 in the DRG. In this study, Shp1-Tg mice appeared to be protected from capsaicin-induced mechanical allodynia, while acute nocifensive behavior and swelling developed similarly in both genotypes. It is possible that in Shp1-transgenic mice, *Shp1* overexpression did not affect all modalities of the capsaicin-sensitive sensory neurons equally. For example, immediate capsaicin-induced TRPV1 activation might be unaffected, leading to acute pain sensation (and nocifensive behavior) and neurogenic inflammation (redness, swelling). However, a genetically enhanced SHP-1 activity and increased dephosphorylation of TRPV1 could diminish TRPV1 sensitization and protect from capsaicin-induced mechanical allodynia. Our interesting finding might suggest a novel strategy for TRPV1 modulation by dephosphorylation, which might overcome the adverse effects of TRPV1 antagonists in the future.

Strengths of our study include the investigation of capsaicin-induced nocifensive behavior in both sexes using a transgenic mouse strain with genetically enhanced SHP-1 expression. A limitation of our study is that thermal hypersensitivity was not investigated due to limited numbers of transgenic mice. Future experiments will explore basal and capsaicin-induced thermal hypersensitivity in WT in Shp1-Tg mice. Also, since SHP-1 has more than one target in sensory afferents, we cannot exclude the effect of SHP-1 overexpression on other substrates such as TrkA, the receptor for NGF. Another limitation is that the investigator was not blinded to the *in vivo* experimental groups, due to the obvious swelling and redness of the paw following capsaicin footpad injection.

We conclude that systemic SHP-1/*Ptpn6* overexpression leads to decreased capsaicin-induced mechanical allodynia, by a mechanism involving TRPV1 dephosphorylation by SHP-1. Our results warrant further investigation into the potential benefit of TRPV1 modulation by SHP-1 on pain related to TRPV1 sensitization and in disease-specific models such as murine models of osteoarthritis.

## Acknowledgements

This work was supported by the following awards: Pilot and Feasibility Grant Award 2022 (PI: AM), provided by the Chicago Center on Musculoskeletal Pain (NIH NIAMS P30AR079206, PI: AMM); Cohn Fellowship (PI: AM) by the Rush Research Mentoring Program, R01AR064251 (PI: AMM), R01AR060364 (PI: AMM) and R01AR077019 (PI: REM) by the National Institutes of Health; Research Foundation Flanders (FWO) Belgium [1842323N and G041519N] and Ghent University [GOA019-21] (PI: FM).

## Author contributions

RV: Data acquisition, data analysis, manuscript editing

SI: Data acquisition

SF: Data acquisition

MW: Data acquisition, data analysis

NA: Data acquisition

NL: Data acquisition

FM: Manuscript editing

AMM: Conceptualization, manuscript editing

REM: Conceptualization, data analysis, manuscript editing

AM: Conceptualization, data acquisition, data analysis, manuscript drafting and editing

## Abbreviations

DMEM: Dulbecco’s Modified Eagle Medium
DMSO: Dimethyl sulfoxide
DRG: Dorsal root ganglia
FBS: Fetal bovine serum
HBSS: Hank’s Balanced Salt Solution
NGF: Nerve growth factor
PD-L1: Programmed death ligand 1
PKA: Protein kinase A
Ptpn6: Protein Tyrosine Phosphatase Non-Receptor Type 6
Scn10a: Sodium Voltage-Gated Channel Alpha Subunit 10
SHP-1: Src homology region 2 domain-containing phosphatase 1
Tg: transgenic
TrkA: Tropomyosin receptor kinase A
TRPV1: Transient receptor potential vanilloid 1
WT: wild type

## Notes

### Competing Interest Statement

A.M.M. is a consultant for Orion, Averitas and 23andMe.

